# Scaffold-level genome assemblies of two parasitoid biocontrol wasps reveal the parthenogenesis mechanism and an associated novel virus

**DOI:** 10.1101/2023.05.25.542369

**Authors:** S.N Inwood, J Skelly, J Guhlin, T Harrop, S Goldson, P.K Dearden

## Abstract

**Background:** Biocontrol is a key technology for the control of pest species. *Microctonus* parasitoid wasps (Hymenoptera: Braconidae) have been released in Aotearoa New Zealand as biocontrol agents, targeting three different pest weevil species. Despite their value as biocontrol agents, no genome assemblies are currently available for these *Microctonus* wasps, limiting investigations into key biological differences between the different species and strains.

**Methods and findings:** Here we present high-quality genomes for *Microctonus hyperodae* and *Microctonus aethiopoides*, assembled with short read sequencing and Hi-C scaffolding. These assemblies have total lengths of 106.7 Mb for *M. hyperodae* and 129.2 Mb for *M. aethiopoides*, with scaffold N50 values of 9 Mb and 23 Mb respectively. With these assemblies we investigated differences in reproductive mechanisms, and association with viruses between *Microctonus* wasps. Meiosis-specific genes are conserved in asexual *Microctonus*, with *in-situ* hybridisation validating expression of one of these genes in the ovaries of asexual *Microctonus aethiopoides*. This implies asexual reproduction in these *Microctonus* wasps involves meiosis, with the potential for sexual reproduction maintained. Investigation of viral gene content revealed candidate genes that may be involved in virus-like particle production in *M. aethiopoides*, as well as a novel virus infecting *M. hyperodae*, for which a complete genome was assembled.

**Conclusion and significance:** These are the first published genomes for *Microctonus* wasps used for biocontrol in Aotearoa New Zealand, which will be valuable resources for continued investigation and monitoring of these biocontrol systems. Understanding the biology underpinning *Microctonus* biocontrol is crucial if we are to maintain its efficacy, or in the case of *M. hyperodae* to understand what may have influenced the significant decline of biocontrol efficacy. The potential for sexual reproduction in asexual *Microctonus* is significant given that empirical modelling suggests this asexual reproduction is likely to have contributed to biocontrol decline. Furthermore the identification of a novel virus in *M. hyperodae* highlights a previously unknown aspect of this biocontrol system, which may contribute to premature mortality of the host pest . These findings have potential to be exploited in future in attempt to increase the effectiveness of *M. hyperodae* biocontrol.

## Background

Species in the genus of *Microtonus* (Wesmael, 1835) (Hymenoptera: Braconidae) have been found to be effective biocontrol agents against forage production weevil pests in Aotearoa New Zealand (NZ), against which they are the primary form of control (Figure 1). *M*. *hyperodae* has been found to provide effective biological control against the Argentine stem weevil (ASW) *Listronotus bonariensis* (Kuschel) (Coleoptera: Curculionidae), a severe pest of Gramineae which causes an estimated NZD$200 million in damage p.a. (Ferguson et al., 2019), while strains of *M. aethiopoides* Loan are effective against the clover root weevil *Sitona obsoletus* (Gmelin) (Coleoptera: Curculionidae), which causes an estimated NZD$235 in damage p.a. (Ferguson et al., 2019), and the lucerne weevil *Sitona discoideus* (Gyllenhal) (Figure 1). behaviours by the weevil (Shields, Wratten, Phillips, et al., 2022; Shields, Wratten, Van Koten, et al., 2022).

**Figure 1.**
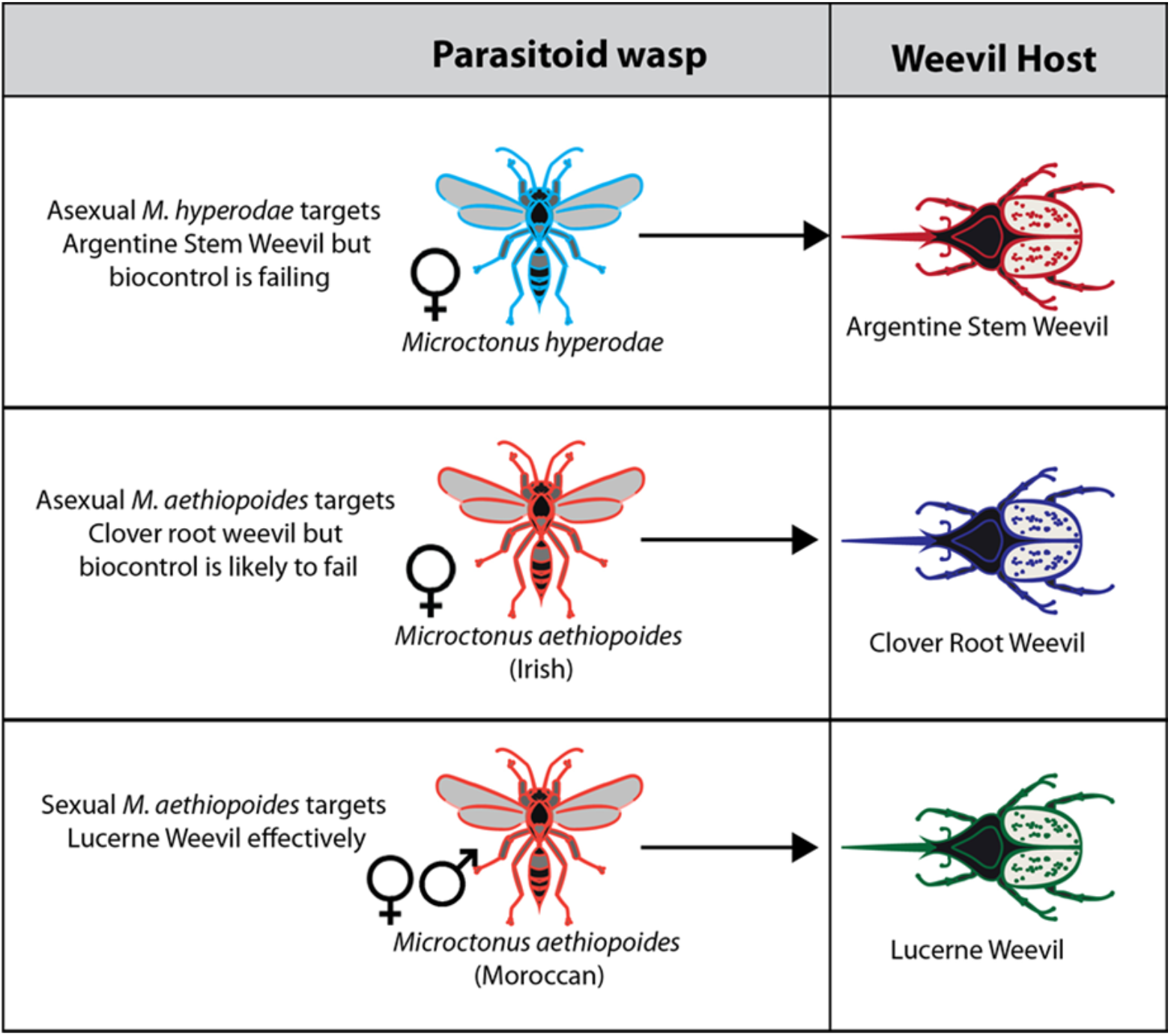
Host target and reproductive mechanisms of *Microctonus* wasps used as biocontrol agents in New Zealand.

*S. discoideus* is controlled by a sexually reproducing strain of *M. aethiopoides* from the Mediterranean area (Stufkens et al., 1987) and is often referred to as the Moroccan strain (e.g. Gerard et al., 2006), released for biocontrol in 1982. This *M. aethiopoides* strain was subsequently found to have no appreciable effect on the invasive populations of *S. obsoletus* (Barratt et al., 1997), resulting in a widespread European search for a suitable parasitoid. A sexually reproducing French strain of *M. aethiopoides* (Goldson, McNeill, et al., 2004) was found to be effective against *S. obsoletus,* however in quarantine it was found to hybridise with the Moroccan strain producing offspring with greatly reduced efficacy against both *Sitona* spp, precluding its use for biocontrol in NZ (Goldson et al., 2003). Continued searching led to the discovery of an asexual European strain of *M. aethiopoides* in Ireland, which reproduces asexually via thelytokous parthenogenesis which would not hybridise with the Moroccan strain (McNeill et al., 2006). This permitted its widespread rearing and release and the eventual suppression of the weevil (Gerard et al., 2006, 2011).

Presence of two reproductive strategies within the *M. aethiopoides* spp. raises interesting questions as to the underlying mechanisms of asexuality in the *Microctonus* genus. While the *M. hyperodae* population in NZ was found to be asexual, four impotent male *M. hyperodae* were discovered amongst the founding 251 the adult *M. hyperodae* reared from weevils from South America (Goldson et al., 1990), indicating some ability to reproduce sexually. Preliminary examination of the biological underpinnings of reproductive mode in *M. hyperodae* using allozymes revealed an absence of recombination, inferred from a lack of certain homozygous genotypes despite prevalent heterozygotes in populations (Iline & Phillips, 2004). From this lack of recombination, it was suggested that *M. hyperodae* parthenogenesis might be apomictic, using mitosis rather than meiosis. However, more recently the conservation and expression of core meiosis genes in *M. hyperodae* ovaries has been demonstrated, indicating that *M. hyperodae* parthenogenesis may involve meiosis thereby retaining the potential for sexual reproduction (Inwood et al., 2023). No such investigation of the parthenogenesis mechanism of the *M. aethiopoides* Irish strain has been performed to date.

Another difference between *Microctonus* wasps is in the presence and transmission of virus particles during parasitism. Endogenous viral elements (EVEs) in the form of polydnaviruses (PDVs) or virus-like particles (VLPs) as well as exogenous viruses, often play a role in the parasitism processes of koinobiont endoparasitoids, particularly in host immune-suppression (Coffman et al., 2022; Di Giovanni et al., 2020; Drezen et al., 2017; Martinez et al., 2012; Ye et al., 2018). Barratt et al. (1999, 2006) detected the presence of viral particles in the ovarian epithelial cells of the sexual Moroccan strain of *M. aethiopoides*, with no viral particles found in other *Microctonus* strains or species. There has been no further investigation into the presence of EVEs or infectious viruses associated with the *Microctonus* wasps, and no investigation using genomic data.

A more comprehensive investigation into the potential presence of viruses or EVEs in *M. hyperodae* is required due to a phenomenon of premature mortality observed in *L. bonariensis,* whereby greater weevil mortality was observed in the presence of an adult *M. hyperodae* than could be explained by parasitism alone (Goldson, McNeill, & Proffitt, 1993; Goldson, Proffitt, et al., 2004; Vereijssen et al., 2011). The current hypothesis is that there may be a toxin-antitoxin system acting during parasitism, whereby *M. hyperodae* transmits something toxic during an unsuccessful ovipositional attempt, which is offset by an ovarian extract during successful parasitism (Vereijssen et al., 2011). This toxin-antitoxin phenomenon has also been observed with the parasitoid *Asobara japonica* Forster (Hymenoptera: Braconidae) and its *Drosophila* host, with the source of toxicity revealed to be virus particles (Furihata et al., 2016; Furihata & Kimura, 2009). The variance in virus particle detection between *Microctonus* species, and the premature mortality phenomenon associated with *M. hyperodae* therefore necessitate an investigation into the viral gene content of the *Microctonus* spp. genome assemblies, and further investigation into the virome of *M. hyperodae*.

Given the range of host targets and reproductive modes, and the importance of biocontrol to a pastoral economy, the genomes of *M. hyperodae* and *M. aethiopoides* have the potential to determine factors that influence biocontrol efficacy, such as genomic correlates of the reproductive mode and host preference, offering a basis for a better understanding the biology of these biocontrol systems. Here we present the scaffolded genomes of two *Microctonus* species, *M. hyperodae* and *M. aethiopoides*, with additional assemblies for a further two *M. aethiopoides* strains with divergent biology, and that of a previously undetected virus found to be infecting *M. hyperodae*, which provide a unique insight into parasitoid reproduction and commensal viruses that play key roles in parasitic life history.

These are valuable genomic resources for understanding the biology of *Microctonus* biocontrol and ongoing investigations into its success or decline in NZ, which is crucial if we are to maintain its efficacy.

## Results and Discussion

### Wasp Genome assembly

Bacterial endosymbionts, particularly *Rickettsia, Wolbachia*, and *Cardinium* can induce parthenogenesis in insects (Ma & Schwander, 2017), while maintaining the potential for sexual reproduction (Arakaki et al., 2000; Stouthamer et al., 1990). As *M. hyperodae* and the Irish strain of *M. aethiopoides* reproduce asexually, the presence of such endosymbionts was investigated using Kraken2 read classification. Kraken2 analysis resulted in the classification of 17.7%, 17.3%, 17.4% and 7.25% of reads from *M. aethiopoides* Irish, French, and Moroccan strains, and *M. hyperodae*, with most remaining unclassified (as the database does not contain insects). Classification of reads as known parthenogenesis-inducing endosymbiont genera was low, with 0.05% or less as *Rickettsia*, 0.01% or less of reads classified as *Wolbachia*, and 0.00% as *Cardinium*. There was no correlation between the percentage of reads assigned to the three genera and the reproductive mechanism of the parasitoids. These results are consistent with previous RNA-seq read classification and PCR results from *M. hyperodae* (Inwood et al., 2023), and with antibiotic and heat treatment failing to revert the asexual reproduction mechanism of *Microctonus* wasps (Phillips, 1995).

Using short-read Illumina sequencing, draft genomes were produced for M. hyperodae, and the Irish, French and Moroccan strains of *M. aethiopoides*, containing 105-128 Mb of total sequence, with a BUSCO completeness of 86.8-93.2% (Table 1). Hi-C scaffolding of the *M. hyperodae* and Irish *M. aethiopoides* assemblies improved these assemblies substantially, with N50 lengths increasing from 15 Kb to 9 Mb for *M. hyperodae* and from 64 Kb to 23 Mb for *M. aethiopoides* Irish (Table 1). The contiguity of these scaffolded *Microctonus* assemblies (Table 1) are comparable to the model insects *D. melanogaster*, *A. mellifera* and *N. vitripennis*, which have N50 lengths of 25.2 Mb, 13.6 Mb and 24.7 Mb, and L50 values of 3, 5 and 7. Compared to other scaffolded Hymenopteran genomes available on NCBI (https://www.ncbi.nlm.nih.gov/data-hub/genome/?taxon=7399, accessed 14/09/22, excluding contig assemblies) the *M. aethiopoides* Irish Hi-C assembly has an N50 length higher than 90% of assemblies, and *M. hyperodae* higher than 75% of assemblies, indicating that these genomes are more contiguous than most scaffolded Hymenopteran genomes, with a high level of BUSCO completeness (Table 1).

**Table 1.** Assembly and annotation statistics for Meraculous and Hi-C scaffolded assemblies for *M. hyperodae* and *M. aethiopoides* strains. BUSCO completeness (%) refers to the number of BUSCO genes found complete (whether single-copy or duplicated) in the genome assemblies.

|  | Hi-C assemblies |  | Meraculous assemblies |  |  |  |
| --- | --- | --- | --- | --- | --- | --- |
|  | M.<br>hyperod<br>ae | M.<br>aethiopoide<br>s Irish | M.<br>hyperoda<br>e | M.<br>aethiopoid<br>es Irish | M.<br>aethiopoid<br>es French | M.<br>aethiopoid<br>es<br>Moroccan |
| Total length<br>(Mb) | 106.7 | 129.2 | 105.8 | 128.8 | 118.9 | 120.3 |
| Number of<br>scaffolds | 3663 | 2844 | 14207 | 6322 | 10395 | 13735 |
| Scaffold N50<br>(bp) | 9364176 | 23025277 | 15392 | 64081 | 30183 | 17930 |
| L50 | 5 | 3 | 1758 | 484 | 881 | 1634 |
| GC (%) | 29.5 | 29.4 | 29.5 | 29.4 | 29.3 | 29.4 |
| Genome<br>BUSCO<br>completeness (%) | 89.8 | 93.9 | 86.8 | 93.2 | 90.7 | 89.1 |
| Predicted<br>genes | 11744 | 12474 | 13667 | 13521 | 14229 | 14475 |
| Annotation<br>BUSCO<br>completeness (%) | 87.9 | 94.0 | 85.2 | 90.7 | 88.4 | 85.8 |

The Hi-C data suggests variance of chromosome number between *M. hyperodae* and *M. aethiopoides*, with 12 and eight Hi-C scaffolds respectively. Investigation of chromosome synteny suggests that chromosomes 1-4 in *M. aethiopoides* are each represented by two chromosomes in *M. hyperodae* (Figure 2), which may represent fusion of chromosomes in *M. aethiopoides*, or fragmentation of chromosomes in *M. hyperodae* in the time since they have diverged.

**Figure 2.**
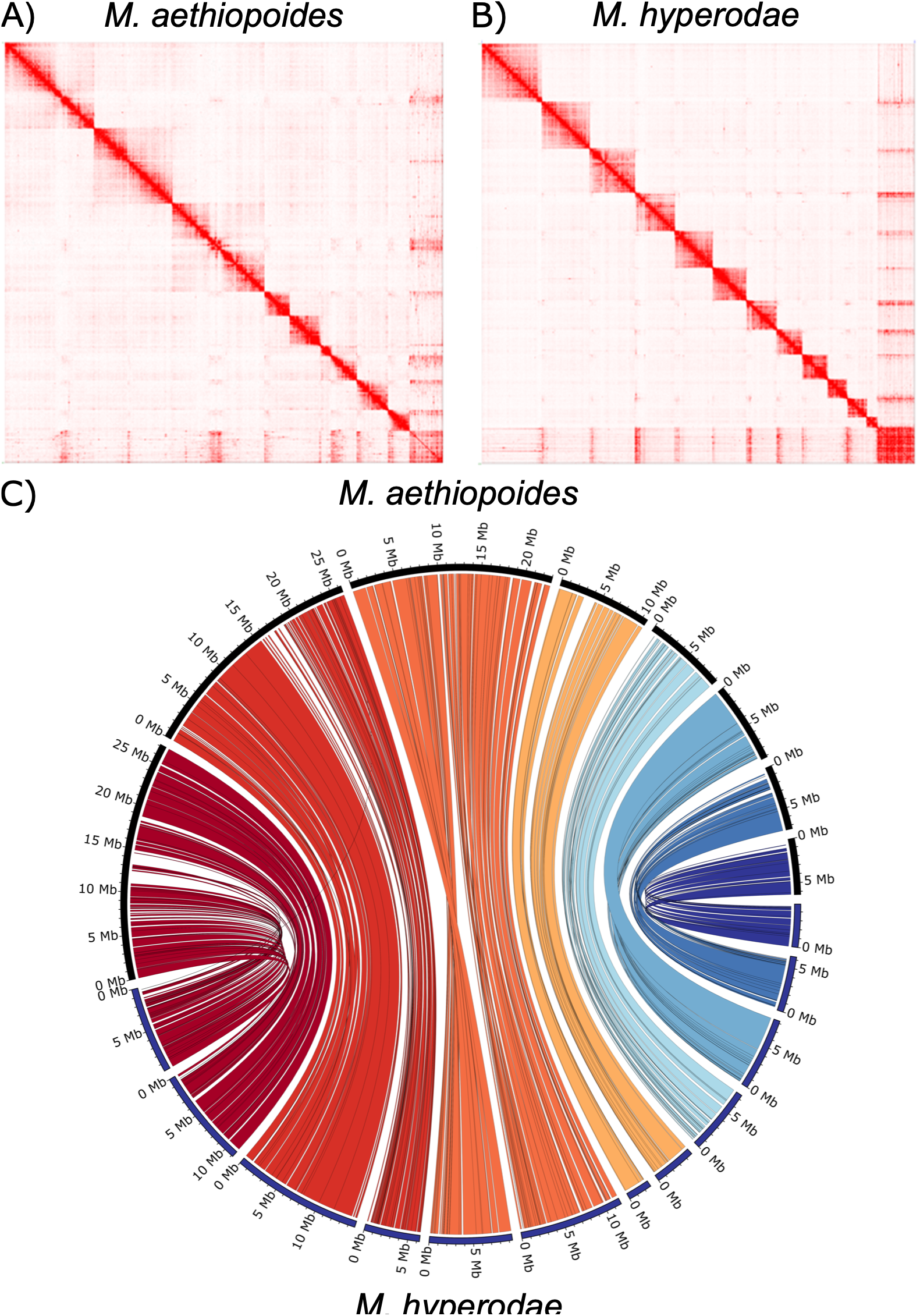
Hi-C interaction maps for A) *M. aethiopoides* Irish and B) *M. hyperodae* displaying chromosome scaffolds, and C) a Circos plot displaying chromosomal synteny between both Hi-C assemblies.

### Wasp Genome annotation

Gene prediction of *Microctonus* genomes was performed with Funannotate, resulting in 12,982 to 14,475 gene predictions for each assembly, with a BUSCO completeness of 85.2 to 94.1% (Table 1). The high gene prediction BUSCO completeness is comparable to other parasitoid wasps (Dalla Benetta et al., 2020; Gauthier et al., 2021) implying that these are high-quality genomes and gene predictions. While Hi-C scaffolding improved assembly contiguity, there was little difference in BUSCO gene prediction results, indicating the short-read only assemblies are still contiguous enough for gene prediction. Only the gene prediction performed for *M. hyperodae* used RNA-seq data (two ovarian samples), and while this may have resulted in improved gene prediction for the *M. hyperodae* assembly, particularly for genes with high expression in the ovaries, the *M. hyperodae* gene predictions, for both the unscaffolded and scaffolded assemblies, had lower BUSCO completeness scores than those for the *M. aethiopoides* assemblies.

Using these high-quality annotations, the divergence between *M. hyperodae* and *M. aethiopoides* strains was estimated, indicating that the French and Moroccan strains (both sexually reproducing) are more closely related than either is to the asexual Irish strain, with the Irish strain estimated to have diverged from the sexually reproducing strains 2 million years ago (MYA), and the two sexually reproducing strains diverging 1 MYA (Supplementary figure 1). The divergence between *M. hyperodae* and *M. aethiopoides* is estimated to be 17 MYA (Supplementary figure 1). This is a larger divergence estimate than was suggested by previous morphological and genetic analysis (Vink et al., 2003).

### Meiosis gene inventory in sexual and asexual strains

A ‘meiosis detection toolkit’ has been described for investigation of asexual reproduction mechanisms in Hymenoptera, cataloguing the presence or absence of genes with known roles in mitosis or meiosis (Schurko & Logsdon, 2008). In an organism that does not use meiosis for gamete production there is no evolutionary constraint on core meiosis genes, so it is expected that their sequence and function would not be conserved (Schurko et al., 2010; Schurko & Logsdon, 2008). Loss of these core meiosis genes would lead to obligate parthenogenesis, where reversion to sexual reproduction is not be possible. This toolkit was used to clarify whether asexual reproduction in *M. hyperodae* and Irish *M. aethiopoides* is automictic, involving meiosis, or apomictic, relying solely on mitosis for egg production which cannot be reverted to sexual reproduction.

This analysis found 36 of 40 meiosis toolkit genes present in single copies in all *Microctonus aethiopoides* assemblies, with the same 36 found in *M. hyperodae* with the addition of RECQ2 (Figure 3). The presence/absence of these genes in these *Microctonus* species is segregated by phylogenetic relatedness, rather than their reproductive mechanism (Figure 3). All but one of the core meiosis genes were detected in *Microctonus* genomes, with only DMC1 being absent (Figure 3), though DMC1 is also absent in *Drosophila melanogaster* and the majority of Hymenopteran species assayed regardless of their reproductive mechanism, implying it is dispensable in meiosis in these species (Tvedte et al., 2017).

**Figure 3.**
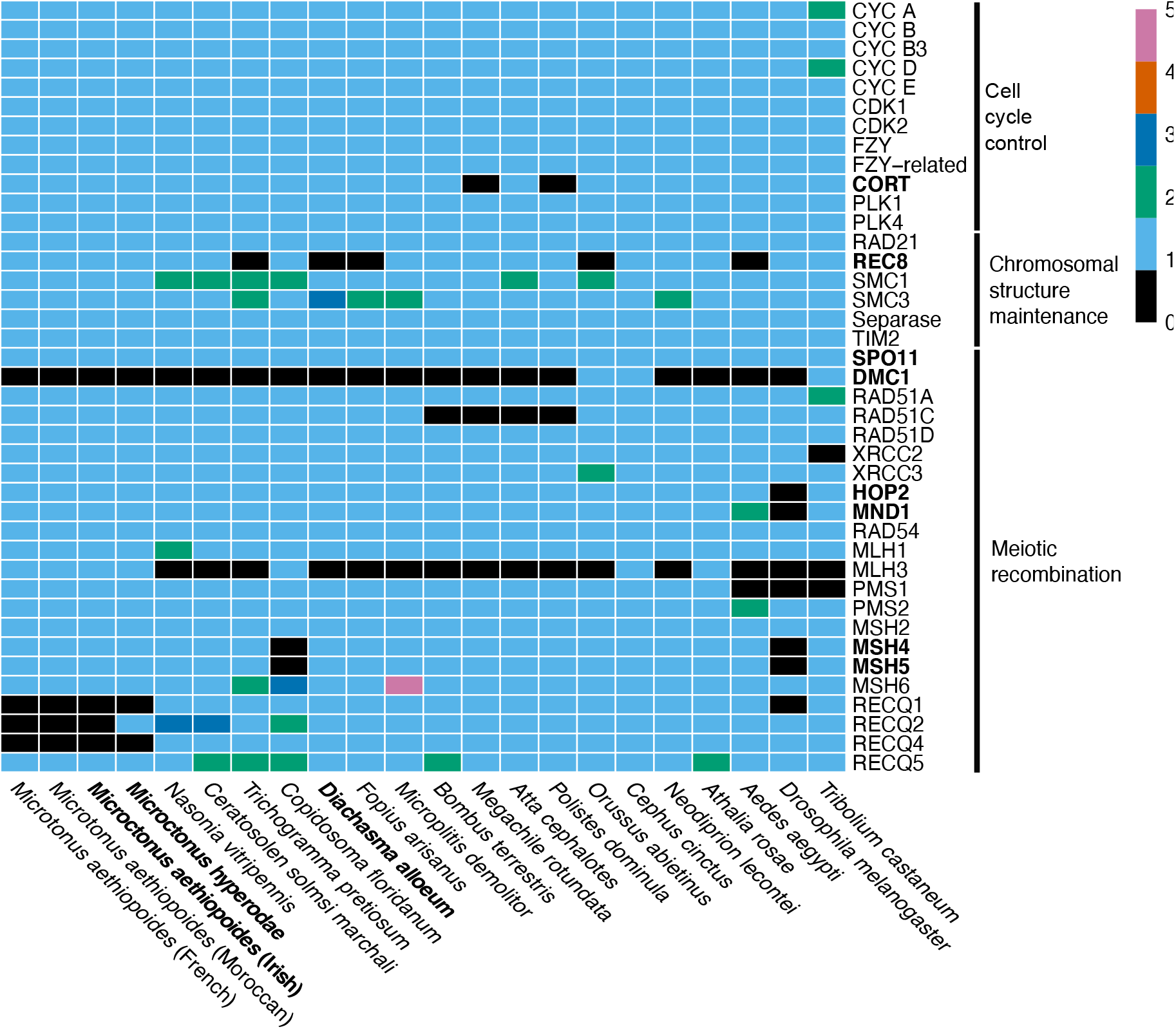
A heatmap displaying the meiosis gene inventory of *Microctonus*, compared to results in other Hymenoptera, Diptera and Coleoptera from Tvedte et al (2017). Colours in heatmap indicate the absence or presence (and gene number) of each meiosis gene. Core meiosis genes, which are specific to meiosis, and asexual Hymenoptera species are indicated in bold.

There is variation in the presence and absence of RECQ genes in the *Microctonus*. RECQ1 and RECQ4 were not identified in the peptide databases of any *Microctonus*, and RECQ2 was only detected in *M. hyperodae* (Figure 3), though this could be due to gene prediction for *M. hyperodae* using RNA-seq data from ovaries while *M. aethiopoides* prediction did not. The RECQ group of proteins are DNA helicases that unwind double-stranded DNA for DNA repair, recombination, and transcription. Some of the RECQ orthologs are required to protect the genome against deleterious mutations (Rezazadeh, 2012). RECQ1 interacts with parts of the DNA mismatch repair pathway during recombination in humans (Doherty et al., 2005).

RECQ4 co-localises with Rad51 after the induction of DNA double-strand breaks, with a possible role in DNA double-strand breakthrough homologous recombination (Petkovic et al., 2005). RECQ2 mutants in *S. cerevisiae* have suppressed non-crossover recombinants indicating a role in mediating recombination product formation (Petkovic et al., 2005). The absence of RECQ genes in *Microctonus* genomes may influence rates of recombination leading to lower genetic diversity, particularly in those that reproduce asexually, and may explain the lack of recombination previously observed in *Microctonus hyperodae* at two allozyme loci (Iline & Phillips, 2004).

Having surveyed the meiosis toolkit genes and examined reads for the presence of endosymbionts that manipulate reproductive mechanisms, we have found no clear evidence for a cause of asexual reproduction in *M. hyperodae* or *M. aethiopoides* Irish. The conservation of most meiosis toolkit genes, particularly all core meiosis genes aside from DMC1 (Figure 3), indicates that asexual reproduction in *M. aethiopoides* Irish and *M. hyperodae* likely uses automixis involving meiosis. This is supported by the detection of MND1 expression in the asexual Irish *M. aethiopoides* ovaries (Supplementary Figure 2), and is consistent with the detection of meiosis core gene expression in *M. hyperodae* ovaries (Inwood et al., 2023), implying that these parasitoids have retained the potential to reproduce sexually.

Heterozygosity rates in these *Microctonus* also do not segregate based on their reproductive mechanisms, with Genomescope Kmer estimates of 0.59%, 0.25%, 0.34% and 0.41% for *M. hyperodae* and the Irish, French, and Moroccan *M. aethiopoides*. The formation of gametes via automixis can involve various strategies to restore diploidy, which have varied effects on rates of heterozygosity, from elimination to retention of all heterozygosity (Pearcy et al., 2006). Previous detection of heterozygosity in *M. hyperodae* (Iline & Phillips, 2004) is supported by these results, with heterozygosity also detected in the asexual Irish *M. aethiopoides*, indicating that if parthenogenesis is automictic in these parasitoids it relies on a strategy that retains at least some heterozygosity. One such mechanism for this is premeiotic duplication which retains all heterozygosity, with observed upregulation of genes involved in endoreduplication in *M. hyperodae* ovaries providing a putative mechanism for this (Inwood et al., 2023). Investigation of where heterozygous sites occur along chromosomes in the Hi-C scaffolded *Microctonus* genome assemblies could assist in determining this mechanism, as was used in investigation of parthenogenesis in the clonal raider ant *Cerapachys biroi* (Oxley et al., 2014).

### Identification of an infectious virus involved in biocontrol

A reciprocal BlastP search of predicted proteins from the *Microctonus* genomes was performed to investigate viral gene content in *Microctonus* genomes, which identified significant viral hits in all assemblies (Supplementary table 2). The number of RNA virus hits was low as expected, given that DNA sequencing should not detect RNA viruses unless their genes are integrated into the genome. A variable number of DNA virus hits were detected in *M. hyperodae* (across 20 contigs) as well in the Moroccan and French strains of *M. aethiopoides* (across five and three contigs) (Supplementary Figure 3). All hits came from viral families known to infect insects, with most *M. aethiopoides* hits to *Polydnaviridae* and most *M. hyperodae* hits to *Baculoviridae* and unclassified DNA viruses (Supplementary Figure 4).

To investigate whether viral genes detected in these assemblies are endogenous (and putatively involved in PDV particle/VLP production) or exogenous, several approaches were taken. First the presence of eukaryotic genes on the same contigs as viral genes was investigated using BlastP and BUSCO. This revealed no strong evidence of eukaryotic genes on viral contigs in *M. hyperodae*, in which no virus-like particles had previously been detected (Barratt, et al., 1999), and recent transcriptome analysis revealed no clear signs of PDVs or other endogenous viral elements with high expression in venom or ovaries (Inwood et al., 2023). *M. hyperodae* was the only parasitoid with a Hi-C genome and DNA virus hit, with none present on the Hi-C scaffolds. This is despite Hi-C data having coverage for the viral contigs, with 0.15% of Hi-C interactions involving viral contigs, and 83.8% of these viral interactions being between two viral contigs, indicating that their exclusion from the main Hi-C scaffolds was not due to these contigs not being sequenced.

Presence of eukaryotic genes on contigs with DNA virus hits was also examined in French and Moroccan *M. aethiopoides*, with no DNA virus hits for the Irish strain. For French *M. aethiopoides* 10 out of 16 genes on these contigs had eukaryotic hits, with all of these belonging to the Braconidae. Similarly, for Moroccan *M. aethiopoides* 11 of 21 genes had eukaryotic hits, seven of which were to Braconidae, providing some evidence that these viral contigs may be endogenous. No BUSCO genes were present on the contigs with DNA virus hits for the Moroccan or French assemblies, which would have provided stronger evidence of whether DNA virus genes are endogenous or otherwise. By contrast, BUSCO genes were present on all contigs with RNA virus hits, indicating they are endogenous as expected. VLPs had previously been detected in the Moroccan *M. aethiopoides*, but not in the limited number of French or Irish samples collected (Barratt et al., 2006; Barratt, Evans, et al., 1999).

The GC sequence content and sequencing depth of viral contigs was compared to Hi-C chromosome scaffolds for *M. hyperodae* and to contigs with BUSCO genes for *M. aethiopoides* strains to provide further evidence of whether viral contigs were exogenous or endogenous (Supplementary Figure 4). There is a significant difference between both the GC content and sequencing depth of viral contigs and Hi-C scaffolds in *M. hyperodae* (GC: mean 34.0% compared to 29.5%, p = 6.7E-08, depth: mean 258 compared to 175, p = 1.6E-03), with this depth result still significant when two high outlier contigs (scaffolds 90 and 995, mean depth of 879) are removed (mean 189 compared to 175, p = 1.3E-02) (Supplementary Figure 4). There was also a significant difference in GC content and depth between BUSCO and viral contigs in *M. aethiopoides* French (GC: mean 35.1% compared to 28.9%, p = 1.9E- 02, depth: mean 172 compared to 281, p = 7.4E- 03) while in *M. aethiopoides* Moroccan there was no significant difference in either metric (GC: mean 32.3% compared to 29.0%, p = 0.12, depth: mean 548 compared to 194, p = 0.88) even when the high depth outlying contig in *M. aethiopoides Moroccan* (depth = 1822), which contains the only *Baculoviridae*, unclassified DNA virus and hemoflagellate parasite hits, was analysed in a separate group (p ≥ 0.50) (Supplementary Figure 4).

Only *M. aethiopoides Moroccan* returned non-significant results for both comparisons (Supplementary Figure 4) providing more evidence supporting that these contigs may be endogenous, with this strain previously demonstrated to have VLPs in their ovaries (Barratt et al., 1999, 2006). No analysis has been done on what these particles contain beyond an inability to extract DNA from them, so it is unknown whether they have a viral origin, though these identified viral genes in *M. aethiopoides* Moroccan are candidates for involvement in VLP production. A more contiguous genome assembly would be required to confirm whether they are exogenous or endogenous. The Hi-C data, GC and sequencing depth analyses do however provide strong evidence that the viral contigs in *M. hyperodae* are derived from an exogenous viral infection.

### *M. hyperodae* metagenome assembly

Given these results, efforts were therefore made to assemble a complete genome for the exogenous virus in *M. hyperodae*. Meta-genomic assembly was performed using MinION long reads, which were generated from a sample that preliminary Illumina sequencing had confirmed infection status with high viral coverage (mean viral depth of 19x, compared to mean Hi-C scaffold depth of 9x). This generated 354,749 MinION reads with a mean 65x depth of the viral contigs and 19x depth of the Hi-C scaffolds. All reads that mapped to the *M. hyperodae* Hi-C scaffolds were removed, retaining 44,388 reads with a mean length of 7305 bp.

Metagenomic assembly from these filtered reads generated 933 contigs, 41 of which were circular assemblies. viralFlye classified one circular contig, six linear contigs and three component contigs with branching paths, as complete viral assemblies. Given the viral BlastP hits to *Baculoviridae* and an unclassified DNA virus, *Leptopilina boulardi filamentous virus* (LbFV), both of which have a circular dsDNA genome, it was expected that the MhFV genome would also be circular. A BlastN search of the complete circular viral contig found significant hits to 18 of the 20 viral contigs (lacking hits to the two high depth contigs excluded from analysis earlier) identified in the *M. hyperodae* genome assembly, indicating that this complete assembly is for the exogenous viral infection identified. Based on the number of BlastP hits to LbFV and later phylogenetic analysis, we provisionally name this virus *Microctonus hyperodae* filamentous virus (MhFV).

### Complete genome assembly of *M. hyperodae* filamentous virus

Illumina read coverage for this complete MhFV assembly was variable (Figure 4), with a maximum of 494x, and a mean of 147x. Illumina short read coverage dropped to 0x in 44 regions, ranging in length from 1 bp to 370 bp, with a mean of 88 bp and total length of 3860 bp. Nanopore read coverage was less variable (with a maximum of 168x and a mean of 69x) and didn’t drop below 4x (Figure 4). The MhFV genome assembly is 163 Kb long, with a GC content of 37.8% and has comparable characteristics compared to other nuclear arthropod-specific large dsDNA viruses (NALDVs) (Supplementary Table 3).

**Figure 4.**
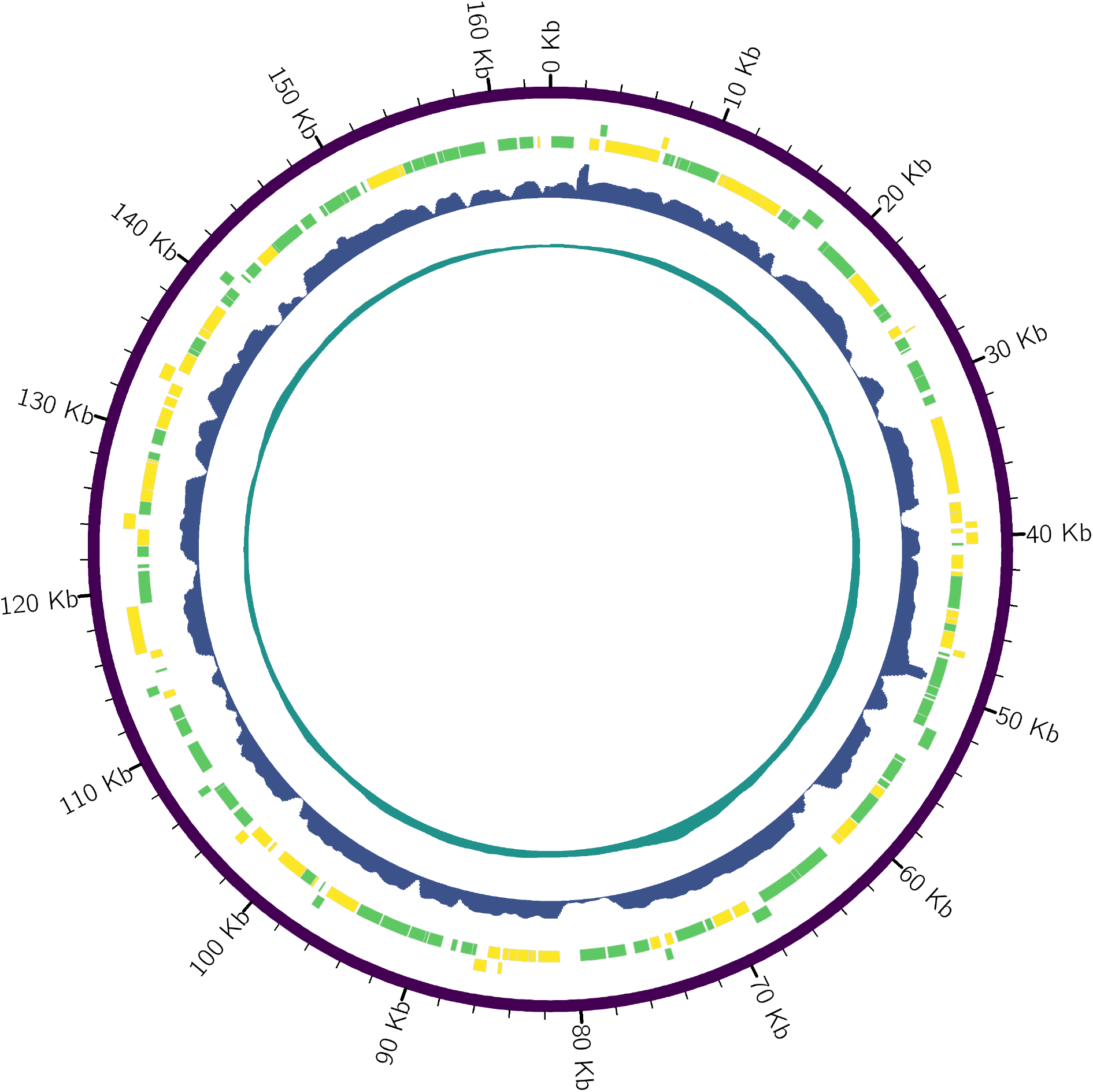
A Circos plot of the MhFV genome assembly. Predicted genes are indicated in the yellow and green blocks, on the positive and negative strand respectively. Relative read depth for Illumina and ONT MinION sequencing is indicated in blue and teal respectively, calculated with a sliding window with a size of 500bp and slide of 100bp.

There are 158 predicted genes in the MhFV genome, all of which are complete and 88.6% of which start with a methionine (minimum amino acid length = 34, maximum = 1652, mean = 286.7). Of the 158 predicted genes, there were significant BlastP hits or Pfam protein domains for 86, leaving 72 genes without any significant homology to indicate potential function (Supplementary Table 4). The most common virus hits were to *Leptopilina boulardi* filamentous virus (LBFV) with 23 hits, followed by *Drosophila*-associated filamentous virus (DaFV, eight hits) and *Glossina pallidipes* salivary gland hypertrophy virus (five hits) (Supplementary Table 4). The predicted genes contain several known viral gene families, such as *per os* infectivity factors, late expression factors, five KilA-N domain-containing genes, and 17 BRO genes (Supplementary Table 4).

While there were 27 hits to bacteria and eukaryotes, an investigation of the full list of Blast hits revealed 11 of these genes also had other viral hits, suggesting potential viral contamination in the assemblies that the best hits came from, rather than there being a putative non-viral origin for all 27 genes. Three genes had only non-viral BlastP hits, including two inhibitor of apoptosis (IAP) genes and a lytic polysaccharide monooxygenase (Supplementary Table 4). ORF32 and ORF152 are both annotated as IAPs (Supplementary Table 4), which are often found in NALDVs and thought to manipulate the host immune system by suppressing apoptosis of infected cells (Crook et al., 1993; Lu & Miller, 1995). IAP genes were also present in the LbFV genome and had a eukaryotic origin (Lepetit et al., 2016). ORF116 and ORF133 are both annotated as lytic polysaccharide monooxygenases (Supplementary Table 4), with at least 97% of hits for both genes from bacterial species.

While none of these hits were viral, many viruses in the families Poxviridae and Baculoviridae contain genes named Fusolin and GP37, which belong to the lytic polysaccharide monooxygenase family (Levasseur et al., 2013). These act to disrupt the peritrophic matrix of insect hosts and facilitate viral infection (Chiu et al., 2015; Phanis et al., 1999). ORF67 was annotated as containing a JmJc-domain (Supplementary Table 4), and while it did have viral hits with one from DaFV and two from different LbFV JmJc-domain containing proteins, there were no other viral hits and 497 non-viral hits. Phylogenetic analysis of the JmJc-domain genes in LbFV indicated they were likely acquired via horizontal transfer from an ancestral host parasitoid (Lepetit et al., 2016). JmJc-containing proteins are a class of demethylase enzymes involved in transcription regulation (Klose et al., 2006).

### Phylogenetic position of MhFV

Recent work identifying endogenous viral elements (EVEs) in parasitoid genomes has identified 12 core genes (*DNApol*, *helicase*, *lef-5*, *lef-8*, *lef-9*, *p33*, *pif-0*, *pif-1*, *pif-2*, *pif- 3*, *pif- 5* and *ac81*) conserved in many NALDVs, and EVEs derived from NALDVs (Burke et al., 2021). Of these core genes, ten were identified in the LbFV genome, missing both *pif-3* and *lef-5*. A BlastP search of gene predictions from the incomplete assembly of DaFV only identified four of the core genes. The MhFV genome has predicted genes with significant BlastP and/or protein domain hits to 11 of these core genes. MhFV is missing only *lef-5*, which is the core gene found in the least number of NALDVs due to its short length (Burke et al., 2021).

Phylogenetic analysis was carried out using protein sequences of the 12 core genes, from all NALDVs used in the analysis by Burke et al., (2021). EVEs derived from NALDVs were excluded, as were the LbFV-like viral contigs identified in that analysis that may have an endogenous origin, as MhFV is an active viral infection rather than an EVE. The resulting phylogeny is globally well resolved, and all characterized viral families were monophyletic displaying expected relationships (Figure 5) (Burke et al., 2021). Despite sharing similar names, *Apis mellifera* filamentous virus and the LbFV-like viruses are separated on the phylogeny, which is the expected result. The placement of LbFV outside of characterized NALDV families is the same result found in other analyses (Burke et al., 2021; Kawato et al., 2019; Lepetit et al., 2016), with the closest family in all previous analyses being

**Figure 5.**
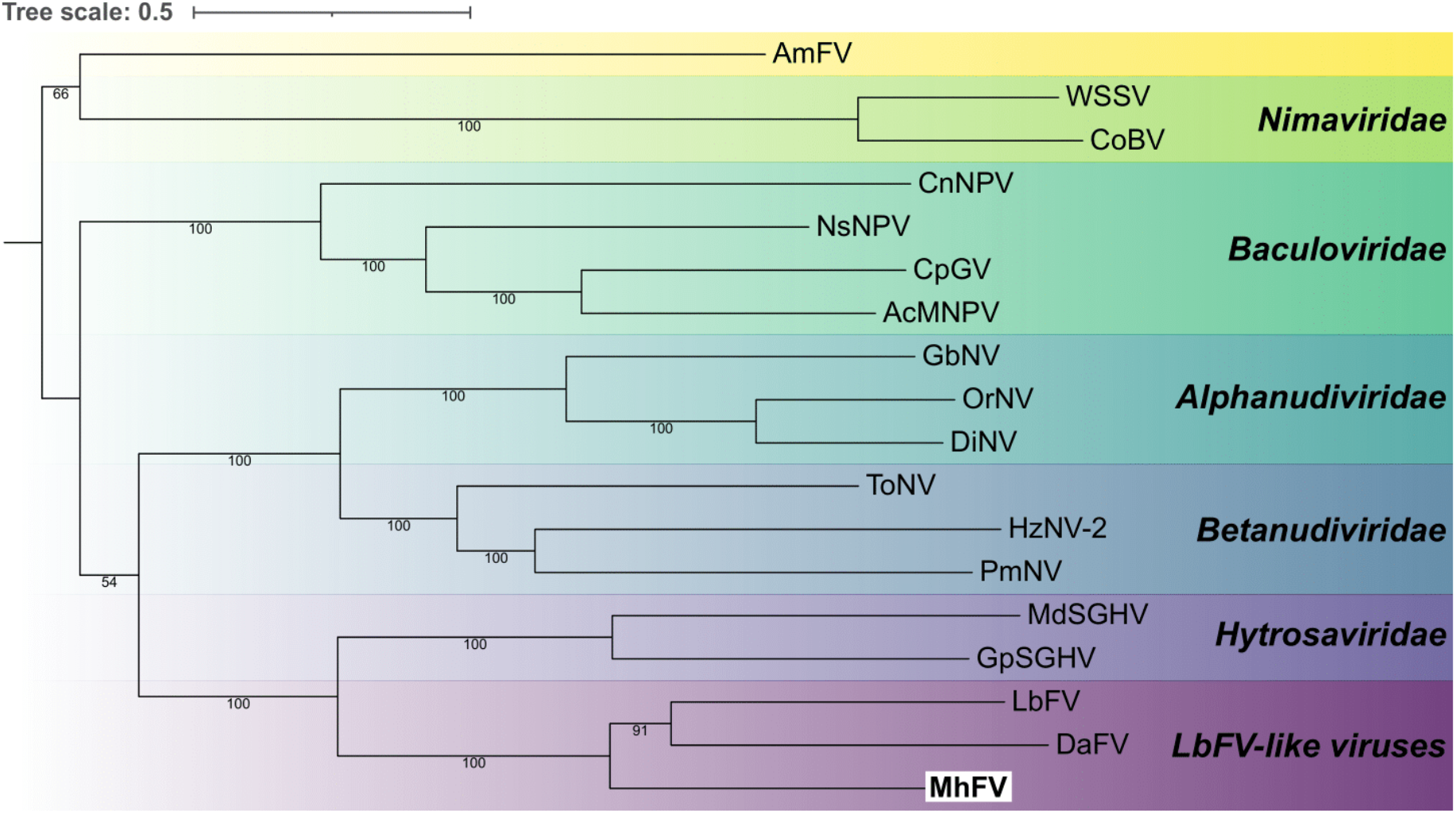
Phylogenetic analysis of nuclear arthropod-specific large double-stranded DNA viruses (NALDVs). Relationships were derived using a maximum likelihood analysis with RAxML-NG, from 12 core NALDV genes, as defined by Burke et al., (2021), with a total of 6818 characters from concatenated amino acid sequences. Bootstrap branch support values over 50% are indicated on relevant branches. Species names are abbreviated as follows; *Apis mellifera* filamentous virus *(AmFV),* White spot syndrome virus *(WSSV), Chionoecetes opilio bacilliform* virus *(CoBV), Culex nigripalpus* nucleopolyhedrovirus *(CnNPV), Neodiprion sertifer* nucleopolyhedrovirus *(NsNPV), Cydia pomonella* granulovirus *(CpGV), Autographa californica* multiple nucleopolyhedrovirus *(AcMNPV), Gryllus bimaculatus* nudivirus *(GbNV), Oryctes rhinoceros* nudivirus *(OrNV), Drosophila innubila* nudivirus *(DiNV), Tipula oleracea* nudivirus *(ToNV), Helicoverpa zea* nudivirus 2 *(HzNV-2), Penaeus monodon* nudivirus *(PmNV), Musca* domestica salivary gland hypertrophy virus *(MdSGHV), Glossina* pallidipes salivary gland hypertrophy virus *(GpSGHV), Leptopilina boulardi* filamentous virus *(LbFV), Drosophila-associated* filamentous virus *(DaFV), and Microctonus hyperodae* filamentous virus *(MhFV)*.

Hytrosaviridae with which a monophyletic group is formed (Figure 5). This analysis places *M. hyperodae* filamentous virus in the same clade as LbFV and DaFV (Figure 5), supporting that MhFV is likely a member of this uncharacterized LbFV-like virus family, though electron microscopy would be required to confirm whether MhFV also has filamentous viral particles.

Manual inspection of mapped Illumina reads revealed the presence of many single nucleotide polymorphisms (SNPs) in the MhFV genome. This variation is likely a result of Illumina sequencing having been performed on a sample with five pooled *M. hyperodae* individuals, as the variability in the MinION reads derived from a single individual is much lower. Variant calling and filtering revealed 1674 SNPs in the MhFV genome, compared to only two in the 111 kb LbFV genome assembly, which was also assembled from a pooled sample (Lepetit et al., 2016). This indicates a relatively high level of variation in the MhFV genome that future population genetics research on *M. hyperodae* should also consider.

### MhFV expression in *M. hyperodae* tissues

A BlastN search of the MhFV gene predictions against a previous *de novo* transcriptome assembly for *M. hyperodae* (Inwood et al., 2023) revealed 79 transcripts with significant hits to 89 MhFV predicted genes. When this RNA-seq dataset is re-examined with the *M. hyperodae* and MhFV genomes, mean TPM values for each predicted MhFV gene in adult tissue samples range from 1.23 (ORF80) to 0.00, while mean TPM in pupa samples ranges from 16.48 (ORF125) to 0.00. The highest mean TPM value for each adult tissue is 3.12 in the abdomen (ORF156), 1.44 in the ovary (ORF88), 1.09 in the thorax (ORF80), 0.64 in the venom gland (ORF125) and 0.31 in the head (ORF156). This data provides transcriptomic support for MhFV infection in *M. hyperodae*, revealing levels of expression are relatively low in adult tissue samples, though this may have been impacted by the use of poly(A) enrichment during library preparation. Differential gene expression analysis revealed 12 MhFV genes with expression patterns significantly influenced by tissue type. These genes had the highest expression in abdomen samples (Supplementary Figure 5), which suggests future efforts to image MhFV particles should focus on tissues in the abdomen.

### Implications of MhFV Infection

MhFV has 23 genes with significant BlastP hits to LbFV, and phylogenetic analysis indicates it is in the same uncharacterized viral family. LbFV infects *Leptopilina boulardi* parasitoids with prevalence varying between 0% to 95% depending on parasitoid density (Patot et al., 2010), and manipulates *L. boulardi* oviposition behaviour, significantly increasing superparasitism of their *Drosophila* host to facilitate horizontal transmission of the virus (Varaldi et al., 2005, 2006). This superparasitism causes decreased overall parasitism rates due to egg wastage, and infected *L. boulardi* are outcompeted by the related parasitoid *L. heterotoma* (Patot et al., 2012). LbFV infection also causes an increase in developmental time and egg load (thought to offset egg wastage from superparasitism), and a decrease in female tibia length.

*L. boulardi* eggs infected with LbFV are also encapsulated significantly less by *Drosophila*, and so this infection may also assist in the avoidance of the host immune system during parasitism (Martinez et al., 2012). This host manipulation could provide another mechanism by which *M. hyperodae* avoids encapsulation of their eggs by *L. bonariensis* (Tomasetto et al., 2017), alongside identified venom components (Inwood et al., 2023), and therefore the potential for MhFV transmission during parasitism should be investigated in future. The possibility for behavioural manipulation of *M. hyperodae* by MhFV should also be considered, but not assumed.

Future research into this biocontrol system should consider the significant impact that the MhFV infection may play in *M. hyperodae* and biocontrol of ASW. MhFV may play a role in the premature mortality phenomenon observed in ASW exposed to *M. hyperodae* (Goldson, et al. 1993; Goldson, et al 2004; Vereijssen et al., 2011), a hypothesis supported by the cause of toxicity during interrupted parasitism attempts with the parasitoid *A. tabida* being identified as viral particles (Furihata et al., 2016). Future work should therefore investigate whether MhFV is transmitted to ASW during successful parasitism and failed parasitism attempts.

Determining the prevalence of MhFV in *M. hyperodae* populations around NZ is also of interest, to see if it is varied between locations with different declines in parasitism rates. If populations without MhFV were identified, these could then be used to determine the physiological effects of MhFV infection on *M. hyperodae*. Examining historical *M. hyperodae* samples, reared and stored from lines imported into NZ from South America before release for biocontrol, should provide evidence of whether MhFV was brought into NZ from their home range, as well as allowing for investigation of whether the prevalence and/or viral load differs in the historical and contemporary samples. Finally, given the large number of variants detected in the MhFV genome with pooled sequencing of five individuals, genetic variation in the virus genome should be considered alongside that in the parasitoid genome, both in comparisons between contemporary locations with different rates of biocontrol decline, and compared to historical samples.

## Conclusions

We generated high quality genomes for the parasitoid wasps *M. hyperodae* and *M. aethiopoides*, which are used as biocontrol agents in NZ, and have investigated aspects of their divergent biology. We have shown that core meiosis genes are conserved in both sexual and asexual *Microctonus* genomes, consistent with previous work implying that asexual reproduction is automictic, involving meiosis, with the potential for sexual reproduction retained. By investigating viral gene content in the genomes, we have also identified candidate genes in the *M. aethiopoides* Moroccan genome that could be involved in VLP production, as well as a novel virus infecting *M. hyperodae*, for which a complete genome was assembled. These resources will be invaluable for future work investigating genomic factors that influence success or failure of *Microctonus*-based biocontrol in New Zealand, and for ongoing investigation of the *M. hyperodae* biocontrol decline.

## Methods

### *Microctonus* spp. samples & sequencing

*Microctonus* spp. samples were supplied by AgResearch, Lincoln,consisting of pools of five*M. hyperodae*, five Moroccan and 10 French *M. aethiopoides* frozen in ethanol as well as 23 live Irish *M. aethiopoides*. DNA was extracted from these samples using a DNeasy kit (Qiagen), and prepared for sequencing using the TruSeq DNA PCR-Free platform. Samples were sequenced on an Illumina HiSeq 2500 by the Otago University Genomics Facility (https://www.otago.ac.nz/genomics/index.html), generating 26, 28, 24, and 35 Gb of 250 bp paired-end reads respectively.

### Genome assembly and scaffolding

The BBTools v37.57 suite BBDuk program (http://jgi.doe.gov/data-and-tools/bb-tools/) was used to trim and decontaminate reads, using default settings for adapter and quality trimming, while also removing the last 5 bp from each read. Kraken2 v2.0.7 was used to taxonomically classify reads against the Kraken standard database (downloaded 17th September 2018) (Wood et al., 2019). Kmer counting was performed using KMC v3.1.1 (Kokot et al., 2017) with a kmer length of 21, for Genomescope2 analysis on the web-based interface (Ranallo-Benavidez et al., 2020).

Meraculous v2.2.5 was used for genome assembly (Chapman et al., 2011), trialling a range of Kmer values, with both trimmed reads and reads normalised by BBNorm to a maximum coverage of 50 and minimum of 5. A range of parameters (including N50 and L50, as determined by the BBTools v38.0 stats wrapper), comparison to estimated genome sizes, and BUSCO v5.4.3 analysis (Simão et al., 2015) were used to determine the best parameters for each genome. Final Kmer values for assembly were as follows: 79 for *M. aethiopoides* French and Irish, 71 for *M. aethiopoides* Moroccan, and 41 for *M. hyperodae*. Normalised reads were used for the French and Moroccan assemblies, and non-normalised for French and *M. hyperodae*. Assembly was run in haploid mode for Irish, and diploid mode for French, Moroccan and *M. hyperodae* assemblies.

Draft genomes for *M aethiopoides* Irish and *M. hyperodae* were scaffolded by Phase Genomics using Hi-C data generated from pools of 10 individuals. Phase Genomics Proximo Hi-C 2.0 kit, a commercially available version of the Hi-C protocol (Lieberman-Aiden et al., 2009) was used to generate chromatin conformation capture data. These data were then used with the Phase Genomics Proximo Hi-C genome scaffolding platform to create chromosome scale scaffolds, as described in Bickhart et al. (2017). BUSCO v5.4.3, and the BBTools v38.0 stats wrapper were then run on Hi-C scaffolded assemblies, with exogenous viral contigs (identification detailed below) removed from the *M. hyperodae* assembly. To investigate genome synteny between the *M. hyperodae* and *M. aethiopoides* Hi-C assemblies, MCScanX was used to identify the 2000 best linkage bundles (Wang et al., 2012), which were then visualised using Circos v0.69-9 (Krzywinski et al., 2009).

### Genome Annotations

Gene prediction was performed using Funannotate v1.5.0-12dd8c7 (Palmer & Stajich, 2019). Repeats were identified using RepeatModeler v1.0.11 (github.com/Dfam-consortium/RepeatModeler) and RepeatMasker v1.5.0 (repeatmasker.org/RMDownload.html) via the Funannotate pipeline. The Funannotate model was then trained using Funannotate train on the masked genome assembly using *Microctonus hyperodae* or *Microctonus aethiopoides* sequences depending on the species of interest. Augustus v 3.3.1 was iteratively trained (Stanke et al., 2008) and used as input for Funannotate predict to predict genes, using the Hymenoptera BUSCO and optimized Augustus settings. Funannotate update was subsequently used to upgrade the annotation. Further annotation was performed with Funannotate using InterProScan 5.32-71.0 (Mitchell et al., 2019). To provide additional data for *M. hyperodae* gene prediction, RNA was extracted from two pools of ovaries using a hybrid of Trizol (Ambion) and RNeasy mini kit (Qiagen) methods. RNA was prepared for using the TruSeq Stranded mRNA platform, and sequenced on an Illumina HiSeq 2500 by Otago Genomics Facility (https://www.otago.ac.nz/genomics/index.html) to generate 250 bp paired-end reads. RNA- seq data was not available for *M. aethiopoides* gene prediction.

To estimate the divergence between *Microctonus* species, Orthofinder v2.3.12 (Emms & Kelly, 2015, 2019) was run in multiple sequence alignment mode, using the diamond search engine and maft, on the genomes of 25 Hymenoptera species and *Drosophila melanogaster*, to produce an ultrametric species tree from peptide databases. The divergence of *Acromyrmex echinatior* and *Camponotus floridanus* of 62 MYA (Peters et al., 2017) was used to calibrate the molecular clock, with a branch length sum of 0.1101, which divided by 62 MYA gave a scaling factor of 563.12. Scaling the ultrametric Orthofinder species tree in Figtree resulted in a phylogenetic tree with inferred divergence estimates, which allowed for divergence between *M. hyperodae* and *M. aethiopoides* to be estimated.

### Identification of mitosis and meiosis-related Genes

To investigate the parthenogenesis mechanism of asexual *Microctonus*, 40 genes known to play a role in meiosis and mitosis (including 8 genes with roles specific to meiosis) were collected for 15 Hymenopteran species, as well as *Aedes aegypti*, *D. melanogaster* and *Tribolium castaneum*, from the NCBI database corresponding to accessions used in a previous analysis by Tvedte et al. (2017). Fasta files were made for the genes and ortholog groups the cyclins (CYC A, CYC B, CYC B3, CYC D and CYC E), the cyclin-dependent kinases (CDK1 and CDK2), CDC20 homologs (CORT, CDC20/FZY and CDC20-like/FZY-related), the Polo-like kinases (PLK1 and PLK4), RAD21 and REC8, structural maintenance of the chromosomes orthologs (SMC1 and SMC3), RAD51 orthologs (RAD51A, RAD51C, RAD51D, XRCC2 and XRCC3), heterodimers HOP2 and MND1, MutL orthologs (MLH1, MLH3, PMS1 and PMS2), MutS homologs (MSH2, MSH4, MSH5 and MSH6), RECQ helicase orthologs (RECQ1, RECQ2, RECQ4 and RECQ5), RAD54, SPO11, TIM2, DMC1, and Separase.

Sequences for each gene and ortholog group were aligned using Muscle v3.8.13 and trimmed using TrimAl v1.2 with the strictplus trimming parameters. Gene-specific hidden Markov models (HMMs) were generated using hmmbuild from Hmmer v3.2.1 (Eddy, 2011). These HMMs were used to search the *Microctonus* peptide databases with hmmsearch, using an E-value of 1E-15. Top hits were selected and peptides retrieved using esl-sfetch from Hmmer v3.2.1. Peptide sequences for each gene were aligned using Muscle v3.8.13 and trimmed using TrimAL v1.2 with the “strictplus” parameters. Phylogenetic trees were constructed for individual genes and ortholog groups using rapidnj v2.3.2 (Simonsen et al., 2008) with a bootstrap value of 1000 for genes and 1000000 for ortholog groups.

### Hybridisation chain reaction in *Microctonus aethiopoides* Irish

Parthenogenetic *Microctonus aethiopoides* (Irish) were reared from CRW collected from pasture in Timaru, New Zealand, with ovaries dissected from 10 wasps three days after eclosion. Ovaries were fixed, and hybridisation chain reaction (HCR) performed with an MND1 probe (with fluorophore 488) as per Inwood et al, (2023). Ovaries were stained with DAPI at a 1:1000 concentration, and mounted on slides in glycerol, for imaging on an Olympus BX61 Fluoview FV100 confocal microscope with FV10-ASW 3.0 imaging software.

### Identification of viral genes in *Microctonus* genome assemblies

In order to identify genes with a viral origin in the *Microctonus* genome assemblies, the predicted peptide databases were searched for peptides with significant homology to viral genes using a reciprocal BlastP v2.9.0 search approach. First a BlastX v2.12.0 search was performed (with an E value of 1E-05 to minimise false-positive results) of all transcripts against the nr database downloaded on May 16^th^, 2021, restricted to only viral entries in the database by using TaxonKit 0.8.0 (Shen & Ren, 2021) to produce a list of all viral taxonomy identifiers at a species level or below and restricting the BlastP search to this list using the - taxidlist option. Any gene with a significant viral BlastX result was then used in a subsequent BlastP search against the whole nr database to remove those genes with better non-viral hits. Finally, any remaining genes on contigs with viral hits that did not have a viral hit were also subject to a BlastP search against the whole nr database to identify potentially eukaryotic genes on the viral contigs. The scripts used for this viral gene identification are available on Zenodo at doi: 10.5281/zenodo.7939016 using a Snakemake-managed workflow (Köster & Rahmann, 2012).

GC content and sequencing depth were compared between Hi-C scaffolds and contigs with DNA virus hits for *M. hyperodae*, and between contigs containing BUSCO hits and contigs with DNA virus hits for *M. aethiopoides* French and Moroccan. The GC content for all contigs was calculated using BBStats v38.0. To determine read depth of contigs,reads were trimmed using BBDuk v38.0 using default settings and trimq=15, and then mapped to the appropriate genome assembly using BWA v0.7.15 (Li & Durbin, 2009). SAMtools coverage v1.10-98 (Li et al., 2009) was then used to determine read mean sequencing depth of contigs. To test for statistical differences between viral and Hi-C/BUSCO contigs, a Shapiro-Wilk test was first carried out to determine whether data were normally distributed, which indicated non-parametric tests should be used. A Kruskal-Wallis rank sum test was used to identify whether there was a significant difference between groups, followed by a pairwise Wilcoxon rank sum test. All statistical tests used an alpha threshold of 0.05 as indication of a significant difference. The scripts used for these sequence characteristic comparisons are available on Zenodo at doi: 10.5281/zenodo.7939021 using a Snakemake-managed workflow.

### Viral genome sequencing and assembly

To completely assemble a genome for the virus detected in *M. hyperodae*, some preliminary Illumina sequence data were examined to identify an *M. hyperodae* sample with high viral coverage. A library was prepared from this sample for long-read Nanopore sequencing on a MinION (Oxford Nanopore Technologies, ONT) using whole-genome amplification and genomic DNA by ligation (SQK-LSK110, ONT), and sequenced on a FLO-MIN106 flowcell (ONT). MinION reads were basecalled using Guppy v6.0.0 with the high-accuracy mode, and Porechop v.0.2.4 used to trim adapters off reads (https://github.com/rrwick/Porechop).

Reads were mapped to the *M. hyperodae* assembly using Minimap2 v2.24 (Li, 2018), to remove reads that aligned to the *M. hyperodae* Hi-C scaffolds. The scripts used for the initial base-calling and read filtering are available on Zenodo at doi: 10.5281/zenodo.7939025 using a Snakemake-managed workflow.

Remaining reads were then used for assembly using Flye v2.9 with the metagenome mode (Kolmogorov et al., 2019, 2020), with output then subject to viralFlye v0.2 analysis (Antipov et al., 2022) to identify which contigs represented full viral genomes. A BlastN search of the only complete circular viral genome assembled revealed significant hits to all *M. hyperodae* viral contigs suspected to belong to the same virus, indicating the complete viral genome had been assembled. Medaka v1.7 (https://github.com/nanoporetech/medaka) and nextPolish v1.4.1 (Hu et al., 2020) were used to polish the assembled genome with long then short reads.The filtered MinION and Illumina reads were then mapped back onto the assembled, polished genome using BWA v2.24, and depth of sequencing at each loci determined using Samtools coverage v1.10-98. Gene prediction was performed using Prodigal v2.6.3 (Hyatt et al., 2010) using the metagenome mode. Predicted protein sequences were subject to a BlastP search against the nr database, retaining all significant hits and filtering results for the lowest E-value and highest bit-score. Hmmscan v3.2.1 was used against the Pfam database (Finn et al., 2014) downloaded 3^rd^ February 2020, to identify protein domains in predicted genes, with results filtered to retain domains with an independent E-value below 5e-04. The scripts used for genome assembly and annotation are available on Zenodo at doi: 10.5281/zenodo.7939027 using a Snakemake-managed workflow.

To determine the phylogenetic placement of *M. hyperodae* virus 1, an approach similar to that used by Burke et al., (2021) was used. This work had identified 12 core genes (*DNApol, helicase, lef-5, lef-8, lef-9, p33, pif-0, pif-1, pif-2, pif-3, pif-5 and ac81*) conserved in NALDVs. All accessions for exogenous viruses used in their phylogenetic analysis were used, and annotations from BlastP and HMMscan were used to identify these genes in the assembled *M. hyperodae* virus genome. To include *Drosophila*-associated filamentous virus (DaFV) in this phylogeny, a BlastP search was performed using the gene predictions available to identify DaFV genes to include in the analysis. Genes and their accessions are detailed in supplementary table 1.

Muscle v3.7 was used to align gene sequences separately, and alignments then concatenated with FASconCAT-G v1.05 (Kück & Longo, 2014). The concatenated alignment was trimmed with default parameters and a gap threshold of 0.6 by TrimAl v1.2. A maximum likelihood phylogeny was constructed for the concatenated alignment using RAxML-NG v1.0.2, with a Blosum62 model and 1000 bootstrap replicates. The phylogenetic tree was drawn using Interactive Tree of Life (Letunic & Bork, 2021), rooted at the midpoint, displaying bootstrap values above 50%. The scripts used for these analyses are available on Zenodo at doi: 10.5281/zenodo.7939029 using a Snakemake-managed workflow.

To investigate the presence of single nucleotide polymorphisms (SNPs) in the assembled virus genome, trimmed reads were mapped to the *M. hyperodae* and virus genomes concatenated into a single file, using BWA v2.24. Bam files were sorted using Samtools v1.10.2, duplicate reads marked with GATK v4.3.0 MarkDuplicates (McKenna et al., 2010) read groups added using AddOrReplaceReadGroups from the Picard tool v2.27.5 (https://github.com/broadinstitute/picard), and resulting bam files indexed with samtools v1.10.2. The bam file was subset to retain only the virus genome, before haplotype calling was performed with GATK v4.3.0 HaplotypeCaller, with ploidy set to five (for a haploid genome with five pooled samples), and variant calling then performed with GenotypeGVCFs. Variants were filtered to retain only SNPs which passed GATK hard-filtering metric recommendations of QD < 2, QUAL < 30, SOR > 3, FS > 60, MQ < 40, MQRankSum < −12.5, ReadPosRankSum < −8, with a minor allele frequency of 0.1. The scripts for virus variant calling are available on Zenodo at doi: 10.5281/zenodo.7939033 using a Snakemake-managed workflow.

To investigate gene expression support for the virus infection in *M. hyperodae*, RNA-seq data from a previous publication (Inwood et al., 2023) was used (NCBI SRA accession PRJNA841753). Reads were trimmed and quasi-mapped as per Inwood et al. (2023) against the *M. hyperodae* and virus predicted genes concatenated into one file. DESeq2 v1.34 (Love et al., 2014) was used to create the DESeqDataSet (DDS) object, by importing Salmon output files using tximport v1.22.0 (Soneson et al., 2016) in R v4.1.3 (R Development Core Team, 2020), with size factors estimated on the concatenated DDS file. The DDS file was then split into separate objects for *M. hyperodae* and MhFV, and DESeq2 then used to perform a likelihood ratio test (LRT) on the virus DDS file with the design ∼Flowcell+Tissue, to control for sequencing runs on different flow cells and test for the influence of tissue on viral gene expression. Differentially expressed genes (DEGs) were identified by filtering DESeq2 results with the arbitrary alpha threshold value of 0.05 for all analyses. A heatmap of viral gene expression was generated using VST normalized data with pheatmap v1.0.12 (Kolde, 2019). The scripts used for this analysis are available on Zenodo at doi: 10.5281/zenodo.7939037 using a Snakemake managed workflow.

## Declarations

### Ethics approval and consent to participate

Not applicable.

### Consent for publication

Not applicable.

### Availability of data and materials

Raw Illumina sequence data, genome assemblies and gene predictions for *Microctonus* wasps are available at the National Center for Biotechnology Information (NCBI) Sequence Read Archive (SRA) and Whole Genome Shotgun (WGS) databases, under accession PRJNA930586, with the Hi-C scaffolded assemblies submitted for *M. aethiopoides* Irish and *M. hyperodae*, with the latter having identified MhFV contigs removed prior to submission. The genome and annotation for *Microctonus hyperodae* filamentous virus is available at Genbank with the accession number OQ439926. Scripts used for analyses are available as detailed in the methods section.

### Author’s contributions

SNI; Data generation, bioinformatics, manuscript drafting, JS; Data generation, bioinformatics, manuscript drafting, JG; bioinformatics

TWH; bioinformatics

SG; Sample collection and curation

PKD; Funding, project conception, supervision, manuscript drafting.

## Acknowledgements

The authors would like to thank Mark McNeill for providing samples and Petra Dearden for proofreading this manuscript.

---

**Supplementary Table 1** Accessions for virus genes used to construct nuclear arthropod-specific large double-stranded DNA virus phylogeny.

**Supplementary Table 2** Viral hits in *Microctonus* genomes from the reciprocal BlastP analysis.

**Supplementary Table 3** A comparison of MhFV virus genome characteristics to other nuclear arthropod-specific large double-stranded DNA viruses.

**Supplementary Table 4** BlastP and HMMscan annotations for MhFV ORFs.

**Supplementary Table 5** Significant differentially expressed genes (DEGs) from DESeq2 MhFV tissue LRT analysis.

**Supplementary** Figure 1 Rudimentary estimation of *Microctonus* evolutionary origin based on an ultrametric Orthofinder species tree, using a branch length scaling factor of 563.12. The *Microctonus* clade is highlighted in the blue dashed box. The “Irish”, “French” and “Moroccan” strains of *Microctonus aethiopoides* are Maeth IR, Maeth FR and Maeth MO respectively. *Drosophila melanogaster* is the outgroup. The divergence of *Acromyrmex echinatior* (*) and *Camponotus floridanus* (*) from Peters et al. (2017) was used to calibrate the molecular clock and is indicated by an arrow. The Hymenopteptera abbreviations are as follows: Nvit (*Nasonia vitripennis*), Tpre (*Trichogramma pretiosum*), Vvul (*Vespula vulgaris*), Vpen (*Vespula pensylvanica*), Vger (*Vespula germanica*), Dnov (*Dufourea novaeangliae*), Mrot (*Megachile rotundata*), Ccal (*Ceratina calcarata*), Amel (*Apis mellifera*), Acer (*Apis cerana*), Bter (*Bombus terrestris*), Bimp (*Bombus impatiens*), Hsal (*Harpegnathos saltator*), Cbir (*Cerapachys biroi*), Lhum (*Linepithema humile*), Fexs (*Formica exsecta*), Cflo (*Camponotus floridanus*), Pbar (*Pogonomyrmex barbatus*), Cobs (*Cardiocondyla obscurior*), Veme (*Vollenhovia emeryi*), Mpha (*Monomorium pharaonic*), Sinv (*Solenopsis invicta*), Waur (*Wasmannia auropunctata*), Aech (*Acromyrmex echinator*) and Acep (*Atta cephalotes*).

**Supplementary** Figure 2 HCR in situ hybridisation of MND1 (red) and DAPI staining (cyan) in the ovaries of asexual “Irish” *M. aethiopoides,* with MND1 expression highlighted by the dashed red box.

**Supplementary** Figure 3 Stacked bar graphs showing the number of significant BlastP hits to viral families for predicted proteins from *Microctonus* genome assemblies. Significant BlastP hits were those with E-values below the E-value threshold of 1E-05. Viral hits are divided into two panels, the upper containing hits to DNA viruses, and the lower to RNA viruses. *M. aethiopoides* Irish is not shown in the upper panel due to having no DNA virus hits.

**Supplementary** Figure 4 Scatterplots and boxplots of mean sequencing depth and GC content of contigs in *Microctonus* genomes. Contigs with viral genes are in purple, and contigs with BUSCO orthologs (for *M. aethiopoides* strains) or Hi-C scaffolds (for *M. hyperodae*), in yellow. Reported P-values were determined using a Kruskal-Wallis test, followed by a pairwise Wilcox test. The number of contigs in each group is reported in the GC content boxplot.

**Supplementary** Figure 5 A clustered heatmap showing expression for all MhFV genes, across the pupa, head, thorax, abdomen, ovaries and venom of *M. hyperodae*, normalized by VST. Genes detected as significantly differentially expressed in the tissue LRT analysis are indicated in bold.

