## Supplementary figures and images for "Scaffold-level genome assemblies of two parasitoid biocontrol wasps reveal the parthenogenesis mechanism and an associated novel virus"

### Supplemental Figures

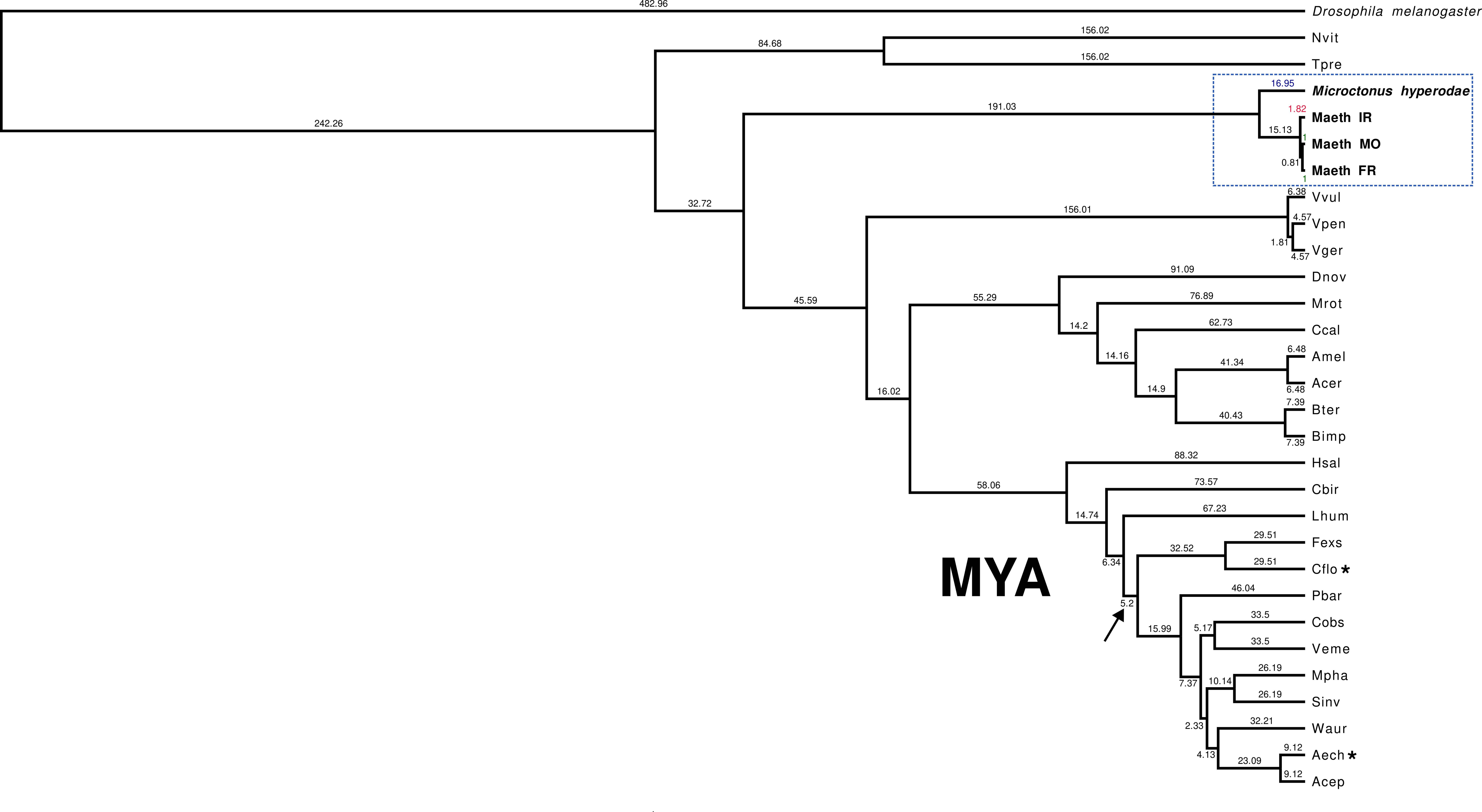

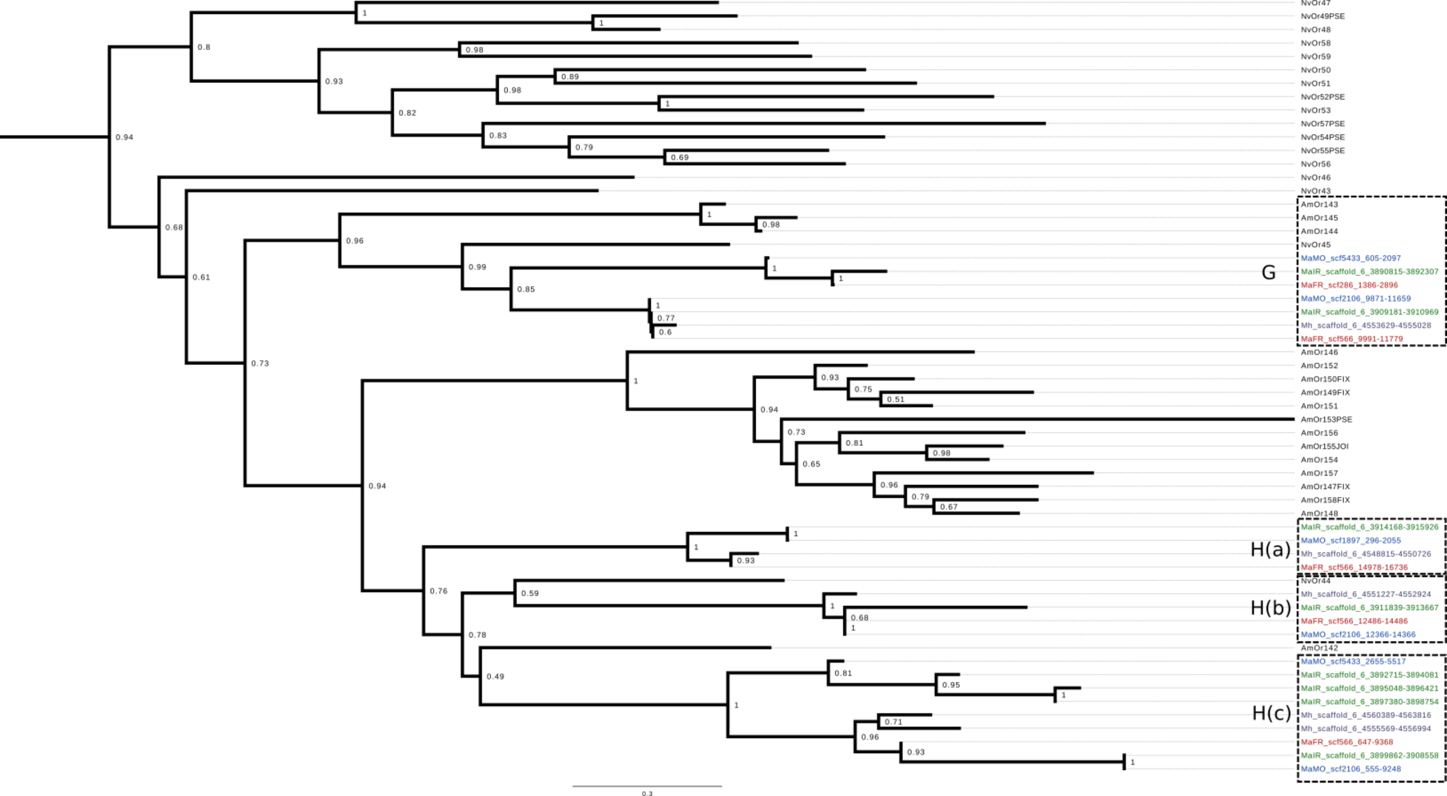

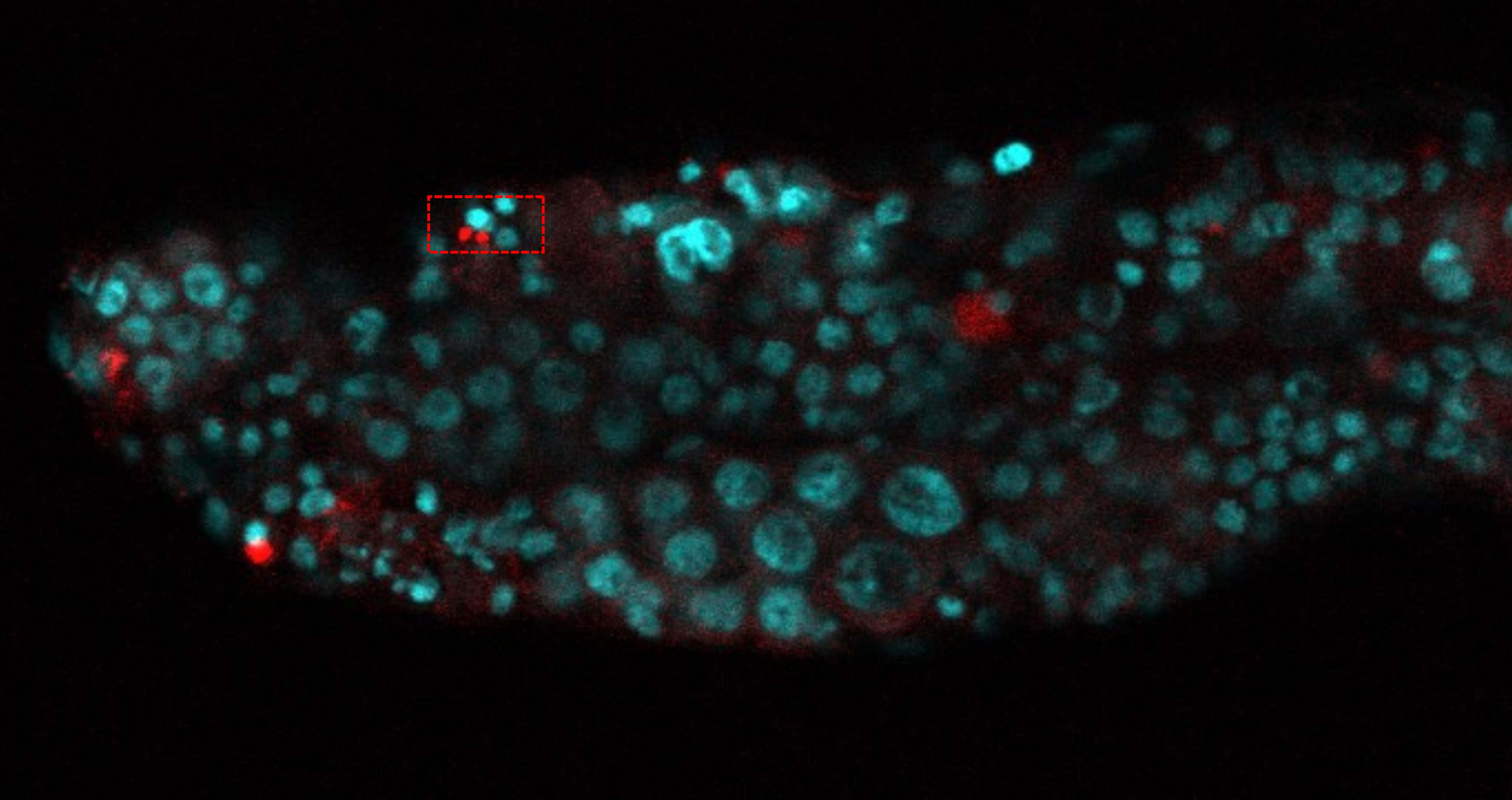

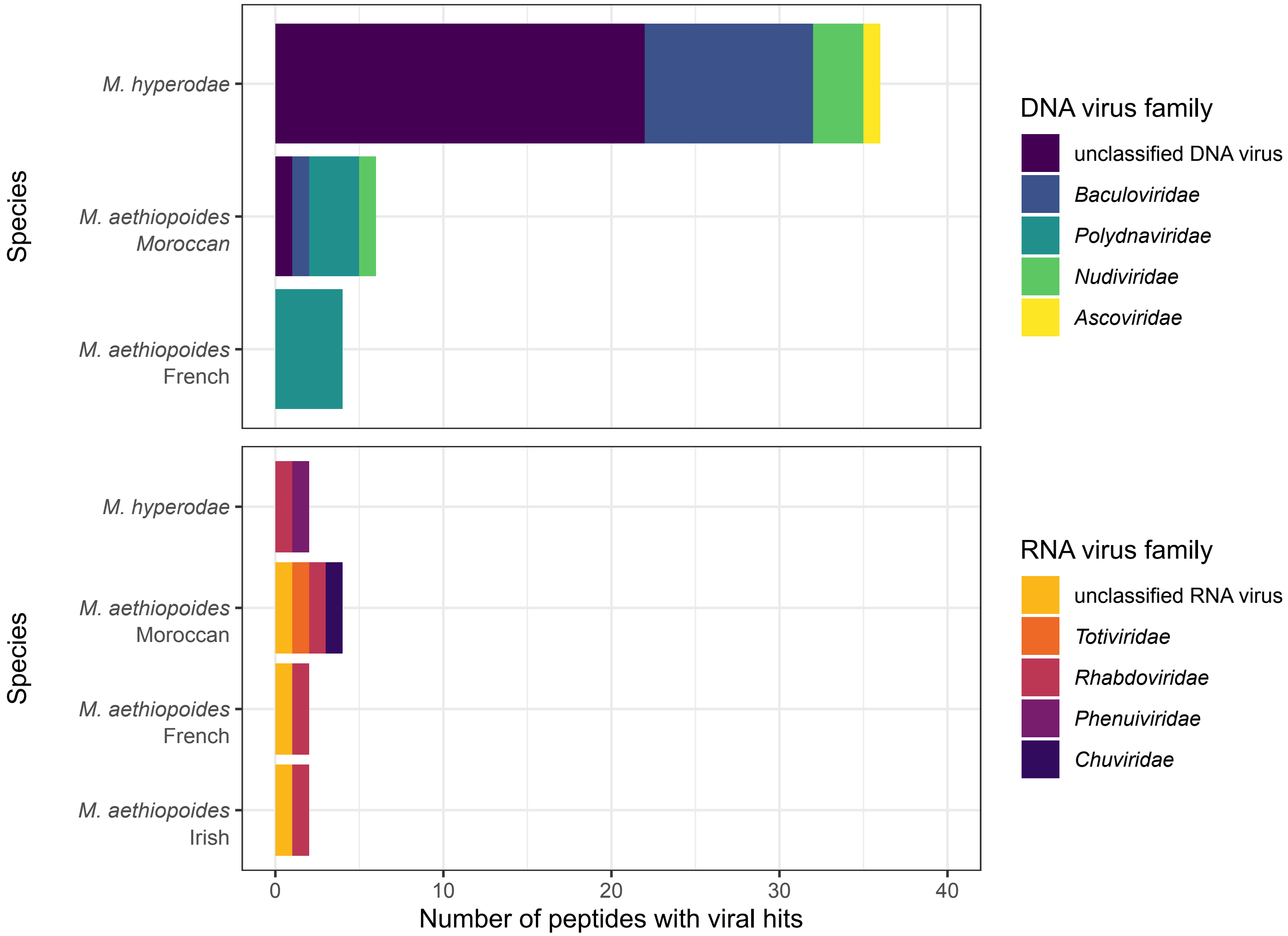

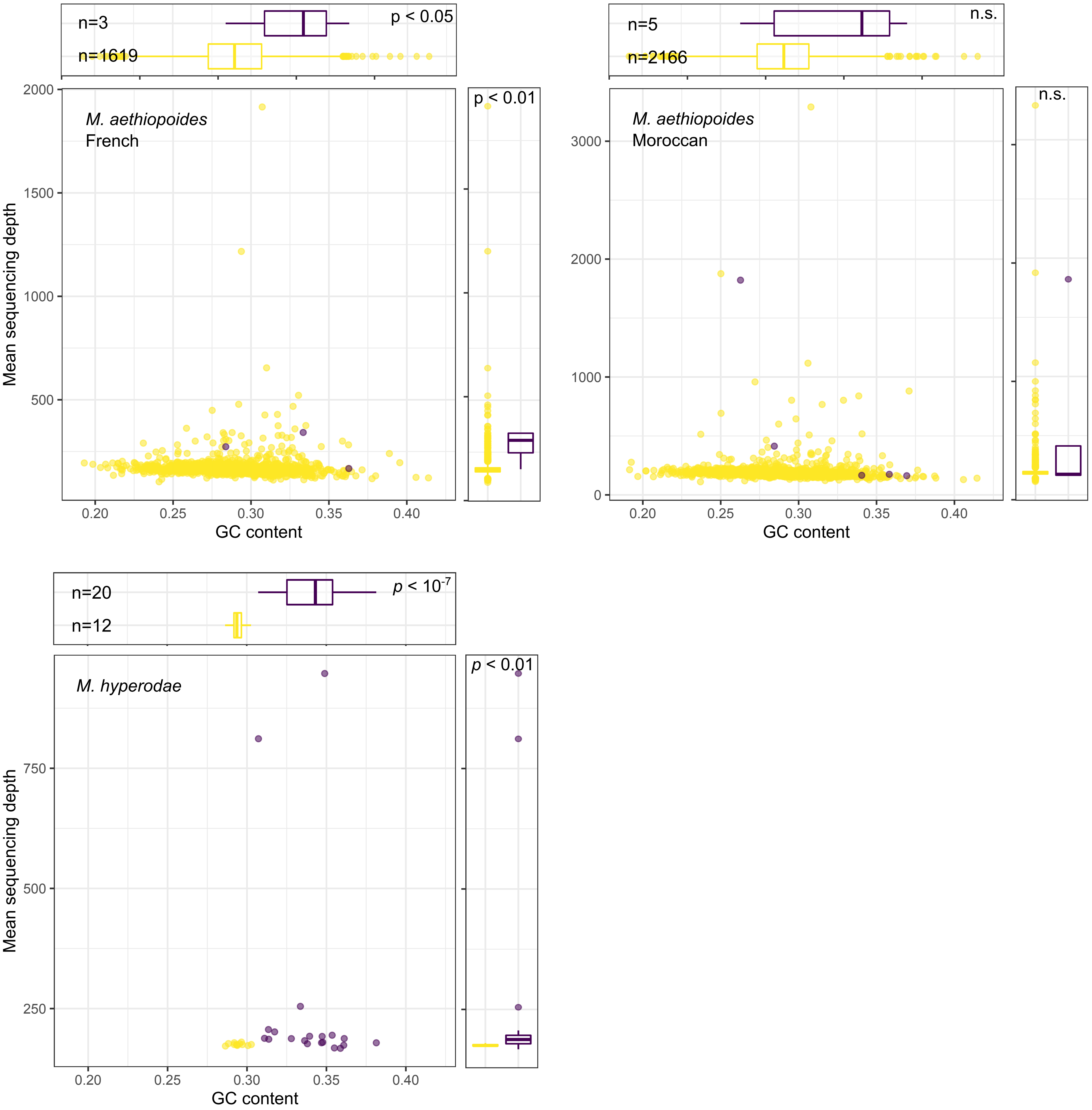

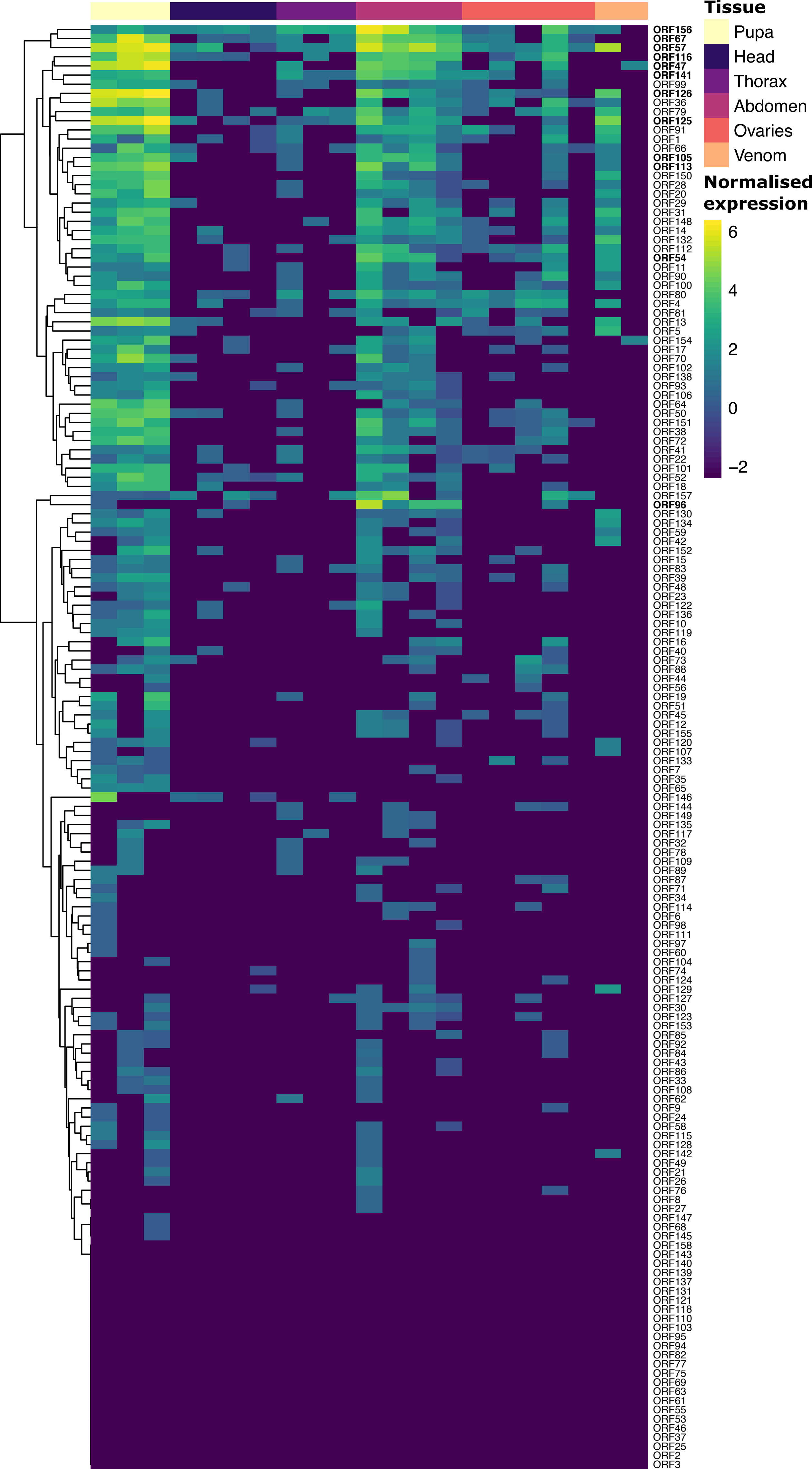
